# Fast-lived Vertebrate Hosts Exhibit Higher Potential for Mosquito-borne Parasite Transmission

**DOI:** 10.1101/2024.10.21.619438

**Authors:** Kyle J.-M. Dahlin, Suzanne M. O’Regan, John Paul Schmidt, Barbara A. Han, John M. Drake

## Abstract

The emergence of mosquito-borne zoonoses has continually increased over the past decade, posing a significant global public health challenge. Ecological theory can point to the characteristics of populations that make them more likely to form reservoirs of disease. The pace of life hypothesis posits that species with more rapid reproduction and shorter lifespans are more likely to be disease reservoirs than their slower-living cousins. Mathematical modeling suggests theoretical conditions under which this hypothesis is correct for directly- and environmentally-transmitted pathogens but its applicability to mosquito-borne disease systems has yet to be examined. We parameterized a mechanistic model with host trait data to investigate the link between the position of a species on the fast-slow life history continuum and the potential for it to spread mosquitoborne diseases. We evaluated the resulting relationships for four medically important mosquito species across their thermal niches and for four pathogens characterized by infection duration and the level of susceptibility of the focal host to infection. Finally, because the fast-slow life history continuum differs across taxonomic ranks, we considered the theory in the context of two orders of vertebrate hosts, Rodentia and Primates. After parameterizing our model, we found that, near universally across all the axes of variation considered, fast-lived hosts have higher transmission potential than slow-lived hosts. There was one exception: slower-lived hosts have higher transmission potential for parasites that cause long-enduring infections to which these hosts are highly susceptible. Generally, however, these connections hinge on the often unknown links between immunological traits, population density, and pace of life. Our analyses highlight immunological traits as a key knowledge gap with strong influence over the potential for transmission. Further investigation of the pace of life hypothesis will require a better understanding of how, for mosquito-borne parasitic infections, susceptibility and infection duration vary across the members of a taxonomic order.

## Introduction

Zoonoses have emerged at greater rates over the past decade (Ndow et al., 2019), a substantial proportion of which are mosquito-borne diseases, such as dengue, chikungunya, and Zika (Bhatt et al., 2013; Gubler, 2002; Messina et al., 2019, 2016). Identifying the animal populations that maintain zoonotic disease transmission can assist with the development of surveillance programs to prevent the spillover of disease into human populations. While the mosquito species which vector certain zoonoses are well-identified, the vertebrate host reservoirs of these same pathogens are often unknown (Althouse et al., 2018; Bueno et al., 2016; Hanley et al., 2013).

Globally, many rodent and non-human primate species are thought to be involved in the transmission of mosquito-borne parasites due to evidence of exposure — through the detection of antibodies in wild populations — or susceptibility and infectivity — as determined by infection experiments on captive populations (Antinori et al., 2021; Sotomayor-Bonilla et al., 2019; Valentine et al., 2019). However, identifying which of these species maintain transmission in their own populations (reservoirs) or contribute to outbreaks in human populations (spillover hosts) remains a significant scientific and public health challenge (Albery et al., 2021; Haydon et al., 2002). Ecological theory can point to the population or life history characteristics of species which make them more likely to function as reservoirs of zoonoses.

In disease ecology, the pace of life hypothesis posits that faster-living host species should be more competent to infection than slower-living species (Huang et al., 2013; Johnson et al., 2012; Ostfeld et al., 2014; Previtali et al., 2012). Mathematical modeling suggests that for directly or environmentally transmitted pathogens, transmission potential is higher for species with higher rates of reproduction and mortality (Han et al., 2020). Similar theoretical work for diseases transmitted by mosquitoes is impeded by the additional layers of ecological complexity involved with vector-borne transmission: the interaction of two types of host and the strong dependence of mosquito life history traits and population dynamics on abiotic factors such as temperature (Mordecai et al., 2019; Thongsripong et al., 2021). Here, we develop this theory as it relates to vector-borne diseases by using compartmental models to evaluate whether transmission potential is higher for vertebrate host species at the “fast” end of the fast-slow life history continuum.

We evaluated the relationship between pace of life and transmission potential in a two-step process. First, using trait data, we determined functional relationships describing the covariation of life history traits for species along a fast-slow continuum. These traits were then used to inform the parameters of a compartmental model of mosquito-borne disease transmission. The basic reproduction number of this model was used to determine the conditions under which an increase in pace of life leads to an increase in transmission potential.

To determine the robustness of our results, we evaluated the relationship between pace of life and transmission potential across several axes of variation. We considered four medically important mosquito species and used empirically-derived thermal performance curves of their traits to incorporate the effect of temperature on transmission. We additionally considered the transmission of four types of pathogens characterized by infection duration in the host and the level of susceptibility of that host to infection. Due to a lack of data on host immunological response to these infections, we examined three degrees of correlation strength between pace of life and infection duration and susceptibility. Finally, because the characterization of the fast-slow life history continuum differs across taxonomic ranks, we looked at two orders of vertebrate host, Rodentia and Primates, which include well-studied species that span both ends of the fast-slow life history continuum.

We find that whether transmission potential is higher for fast-lived vertebrate hosts hinges on the relationships between population density and immunological traits on pace of life. Faster-lived species that form dense populations have higher transmission potential if immunological traits are strongly correlated with pace of life. On the other hand, if faster-lived hosts form more sparse populations, then transmission potential increases along with pace of life only when immunological traits are weakly associated with pace of life.

After parameterization, our model suggests that across mosquito species, temperatures, and pathogen types, faster-lived hosts have higher parasite transmission potential. However, when parasites cause chronic infections and hosts have high susceptibility to infection, slower-lived hosts can have higher transmission potential than fast-lived hosts. But this situation only occurs when the ratio of mosquito to host abundance is suitably small.

Our analyses revealed that detailed information about immunological traits is a key knowledge gap with strong influence over predictions about transmission potential. While our modeling suggests that populations of fast-lived hosts have a greater potential to transmit mosquitoborne parasites, these results are conditioned on assumptions about how immunological traits vary with pace of life, necessitated in our study by the lack of data on these traits. Comparative studies of host immunity would provide valuable insights into what types of animals are most likely to harbor mosquito-borne parasites.

## Methods

### Model

We start with a compartmental model simulating the transmission of a mosquito-borne pathogen between a mosquito population and a vertebrate host population that was originally presented in Dahlin et al. (2024) and is illustrated in Figure 1. This model. Vertebrate hosts are recruited at the per-capita rate *λ*_*H*_, experience natural mortality at the per-capita rate *µ*_*H*_, and have a carrying capacity of *K*_*H*_ individuals. Vertebrate hosts are contacted by mosquitoes at the per-capita rate *b*_*H*_(*H, V*) which may be a function of the total population sizes of hosts *H* and mosquitoes *V*. Susceptible vertebrate hosts may become infected with probability *β*_*H*_ when an infectious mosquito feeds on their blood. After becoming infected, the vertebrate host is, on average, infectious for 1/*γ*_*H*_ days before recovering with permanent immunity. Mosquitoes are recruited at the constant rate *λ*_*V*_ while experiencing mortality at the per-capita rate *µ*_*V*_, so that their carrying capacity is given by *λ*_*V*_ /*µ*_*V*_. Mosquitoes make contact with vertebrate hosts at the per-capita rate *b*_*V*_ (*H, V*) which may also be a function of the total population sizes of hosts (*H*) and mosquitoes (*V*). Similarly, susceptible mosquitoes may become infected with probability *β*_*V*_ after feeding on an infectious vertebrate host. An infected mosquito survives long enough for the pathogen to develop (the extrinsic incubation period) with probability *θ*_*V*_ before becoming infectious and remaining infectious for its lifespan. The full system of equations for the model is:

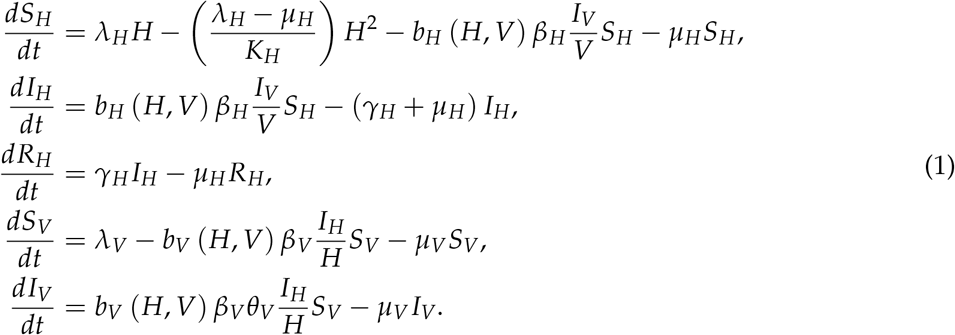

*H* = *S*_*H*_ + *I*_*H*_ + *R*_*H*_ represents the total vertebrate host abundance and *V* = *S*_*V*_ + *I*_*V*_ the total mosquito abundance.

**Figure 1:**
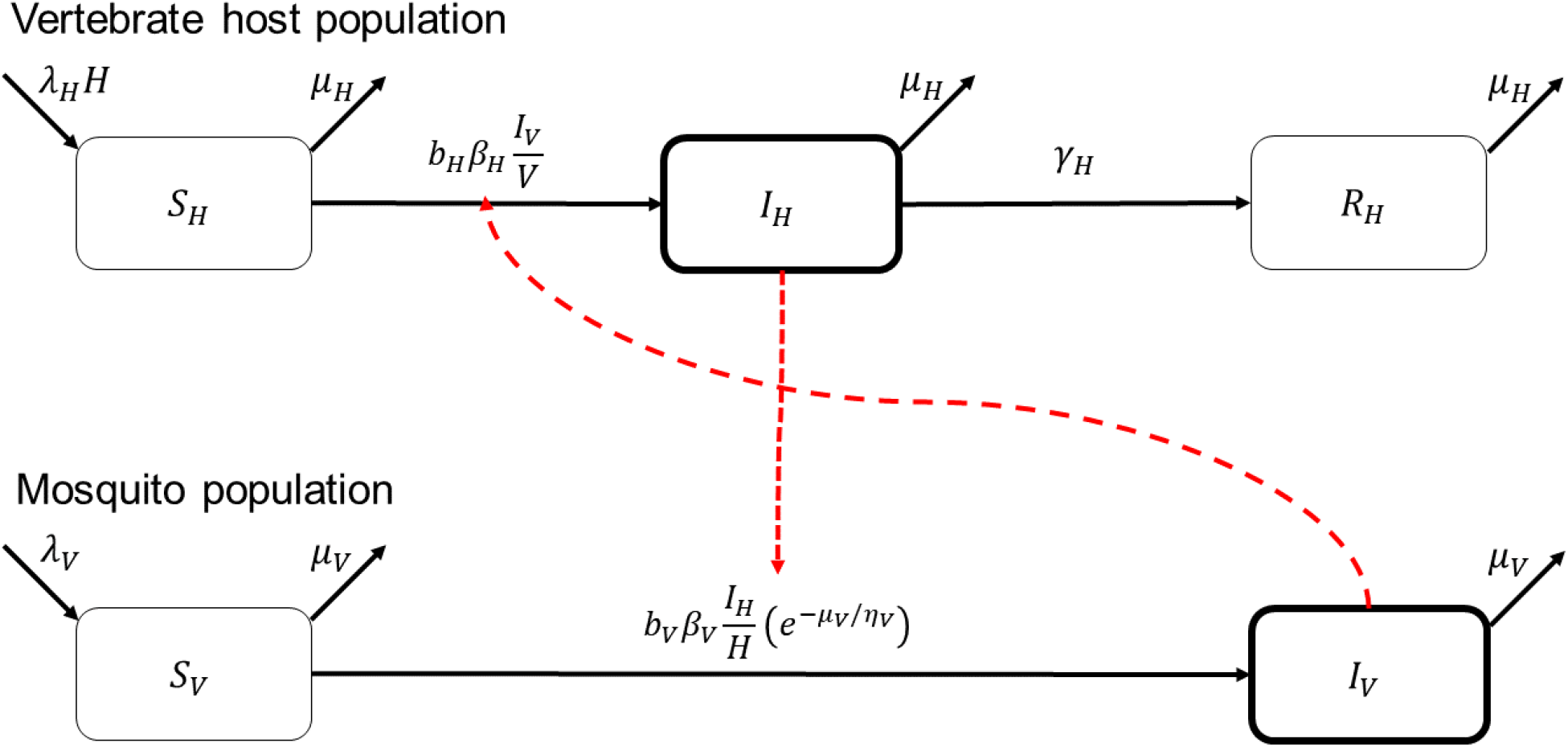
Flow diagram for our compartmental model of mosquito-borne parasite transmission. Black arrows denote transitions of individuals between states while red dotted arrows indicate the influence of one compartment on new infections in another. In this model, *S* denotes susceptible individuals of either hosts (*H*) or vectors (*V*), *I* denotes infectious individuals, and *R* denotes recovered (and immune) individuals.

Because we wanted to consider a range of values for vertebrate host traits, including population density, we considered an alternative to the Ross-Macdonald formulation of the contact rates that does not have an asymptote at low vertebrate host population densities (Chitnis et al., 2006). Modelers of mosquito-borne disease transmission typically assume that either the ratio of vector and host abundances is fixed or, more generally, that incidence is proportional to the vector-host ratio (Reiner et al., 2013). The Chitnis dynamic formulation of contact rates instead bounds the contact rates for large values of the vector-host ratio by incorporating the assumption that there is a maximum value, *σ*_*H*_, to the number of mosquito bites that a vertebrate host would tolerate in a given period of time. In this model, the rates are given by the functions:

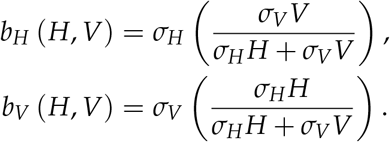

The classical Ross-Macdonald contact rate assumption (as in (Smith et al., 2012)) is a special case of the Chitnis dynamic contact rate formulation where the biting tolerance of the host is instead unlimited, i.e., 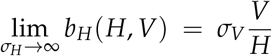 and 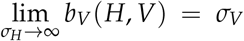. The maximum number of mosquito bites tolerated by a host per unit time (*σ*_*H*_) is an unknown parameter that we assumed is equal to 100 for primates and 10 for rodents. We also considered the case where *σ*_*H*_ → ∞ for both clades.

### Parameterization

#### Quantifying pace of life

The pace of life syndrome is commonly associated with a fast-slow life history continuum characterized by traits with units of time frequency or time duration, such as longevity or fecundity (Van de Walle et al., 2023). Slow-lived species are assumed to reproduce more infrequently and to have longer lifespans while fast-lived species reproduce more rapidly and have shorter lifespans. We aim to derive a parameter *p* that corresponds to a species’ position on the fast-slow life history continuum for a given order of animals.

To facilitate our analysis of the effect of pace of life on transmission potential, we suppose a synthetic variable *p* represents the position of a species in the fast-slow life history continuum with smaller values of *p* corresponding to species with a slower life history. Parameters corresponding to the life history traits of each vertebrate host species were fit as functions of *p* so that the covariation among model parameters best represented the available empirical trait data. There are five host parameters that we assume depend on *p*: recruitment rate (*λ*_*H*_), population density (*K*_*H*_), mortality rate (*µ*_*H*_), the probability of a host becoming infectious given contact with an infected mosquito or susceptibility for short (*β*_*H*_), and recovery rate (*γ*_*H*_).

We used a set of trait data for primates obtained from the PANTHERIA database (results for rodent trait data are qualitatively similar and are given in the Supplementary Material). Missing data for traits were imputed using the missForest algorithm from the R package of the same name (Stekhoven and Bühlmann, 2012). Taxonomic data were not included in the imputation algorithm for the results shown here; including taxonomy did not sufficiently change our results (see Supplementary Material). Reproductive rates were taken to be the product of litter size and litters per day while mortality are the inverse of maximum longevity in days.

In past studies, the position of a species on the fast-slow life history continuum has been quantified through various methods including the generation time of the species (a quantity derived from multiple life history traits) or the first principal component of the traits of the species (Gaillard et al., 2016; Van de Walle et al., 2023). We considered several alternative metrics for pace of life including generation time, maximum lifetime reproductive output, and the loading of the first principal component, among others (see the Supplementary Material for a full description). When it was necessary to decide the direction of the relationship between a metric and life history parameters, the directionality was set so that reproduction rate and mortality should increase with the metric. Each species was ordered according to each pace of life metric and assigned a value so that the “slowest” species was assigned to zero and the “fastest” to one. The model parameters corresponding to life history traits (population density and reproduction and mortality rates) were then fit as exponential functions of *p* using the same primate trait data set. That is, a life history parameter (*z*) is assumed to relate to *p* as *z*(*p*) = *z*_0_*e*^*cp*^ where *z*_0_ is the parameter value at the “slowest” end of the continuum and *c* = log *z*(1)/*z*(0) is a measure of the total amount of variation in the trait across the spectrum. We also fit these parameters as linear functions but this resulted in a poorer explanation of the variance in the traits.

#### Immunological parameters

Traits affecting the immune response of an animal may also be strongly linked to the pace of their life (Martin et al., 2007; Ostfeld et al., 2014; Pap et al., 2015; Previtali et al., 2012; Tieleman, 2018). Despite evidence of these links, the available data is insufficient to directly parameterize the covariation among immunological and life history traits. We therefore considered a range of ways in which immunological and life history traits might covary. We considered four “types” of parasite characterized in two ways: i) the duration of infection (acute or chronic), and ii) the level of susceptibility of the hosts to infection (low or high). The parasite type then parameterizes the relationship between *p* and the recovery rate (*γ*_*H*_) and susceptibility (*β*_*H*_) parameters which, in analogy with the life history model parameters, are also exponential functions. Baseline values for infection durations were chosen to roughly correspond to real parasites. We chose a baseline of two days for acute infections, which, for parasites like dengue virus, can last a few days to a week in non-human primates (Muhammad Azami et al., 2020). Chronic infections were assumed to last for a proportion of the vertebrate host’s lifespan, starting at 1%, which roughly matches the infection duration of parasites like malaria in primates (Ashley and White, 2014; Bretscher et al., 2011). To further expand our analyses, we considered three levels of variation in these immunological traits: no variation, “narrow” variation, and “wide” variation. The wide case features twice as much variation as the narrow case. Table 1 shows the range of values of these immunological parameters as *p* varies from zero to one.

**Table 1:** Ranges of values for immunological traits as pace of life is varied for the four types of parasites and two levels of variation. As pace of life (*p*) is increased from zero to one, parameters increase from their minimum to the maximum values according to the function *z*(*p*) = *z*_0_*e*^*cp*^.

| Trait | Minimum | Maximum |
| --- | --- | --- |
| Life history traits |  |  |
| Reproductive rate, $\lambda_H$ | $1.53 \times 10^{-3}$ per day | $4.20 \times 10^{-3}$ per day |
| Mortality rate, $\mu_H$ | $7.76 \times 10^{-5}$ per day | $1.58 \times 10^{-4}$ per day |
| Population density, $K_H$ | $1.46 \times 10^{-1}$ per ha | $6.35 \times 10^{-1}$ per ha |
| Immunological traits |  |  |
| Susceptibility, $\beta_H$ | | |
| Low baseline, narrow variation | 5% | 25% |
| Low baseline, wide variation | 5% | 45% |
| High baseline, narrow variation | 60% | 80% |
| High baseline, wide variation | 60% | 100% |
| Infection duration, $1/\gamma_H$ | | |
| Acute infection, narrow variation | 2 days | 7 days |
| Acute infection, wide variation | 2 days | 12 days |
| Chronic infection, narrow variation | 1% of lifespan | 3% of lifespan |
| Chronic infection, wide variation | 1% of lifespan | 5% of lifespan |

#### Temperature dependence of mosquito traits

To explore whether the identity of the mosquito and parasite species impacts our findings, we parameterized our model for four medically-important mosquito species. Mosquito traits were assumed to follow thermal performance curves derived from empirical data following (Johnson et al., 2015; Mordecai et al., 2019, 2017, 2013; Shocket et al., 2020; Tesla et al., 2018). Specifically, thermal performance curves were used to determine biting rate (*σ*_*V*_), fecundity (*λ*_*V*_), adult mortality rate (*µ*_*V*_), the probability of surviving the extrinsic incubation period (*θ*_*V*_), and vector competence (*β*_*V*_) as functions of temperature. Because our focus is not on a specific parasite, we assumed that the parasite development rate was independent of temperature and that the average extrinsic incubation period was six days.

### Mathematical analysis

Transmission potential was measured through the basic reproduction number of the model (1), ℛ_0_:

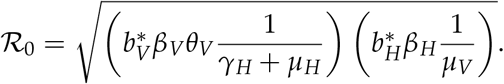

To assess whether there is quantitative evidence for the pace of life hypothesis, we re-interpreted the hypothesis numerically as the property that *R*_0_ increases as a function of *p*, that is, *d*ℛ_0_(*p*)/*dp* > 0. Because it takes a more easily interpretable form, we evaluated the logarithmic derivative of ℛ_0_ as a function of *p*, which we denote 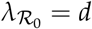 ln ℛ_0_/*dp*, and which has the same sign as the usual derivative. The value of 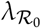 is then calculated across all of the axes of variation previously described — position on the pace of life spectrum, parasite type, mosquito species, and temperature — to determine in which cases 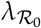 is positive.

## Results

### Quantifying pace of life

Among the possible measures of pace of life considered, generation time, the time it would take the population to double calculated as the ratio of log(2) and the intrinsic population growth rate, best explained the covariation between reproductive and mortality rates (see Supplementary Figure S1) and, after normalizing values to be between zero and one, is hereafter simply denoted by *p*. Figure 2 shows the fitted relationships between the life history parameters and *p* for the rodent and primate trait datasets. While there was no assumption on the relationship between population density and pace of life, each of our measures of pace of life produced a weak but positive relationship between *p* and population density, *K*_*H*_. Table 1 gives the range of values taken by the life history traits as *p* is varied from zero (Minimum) to one (Maximum).

**Figure 2:**
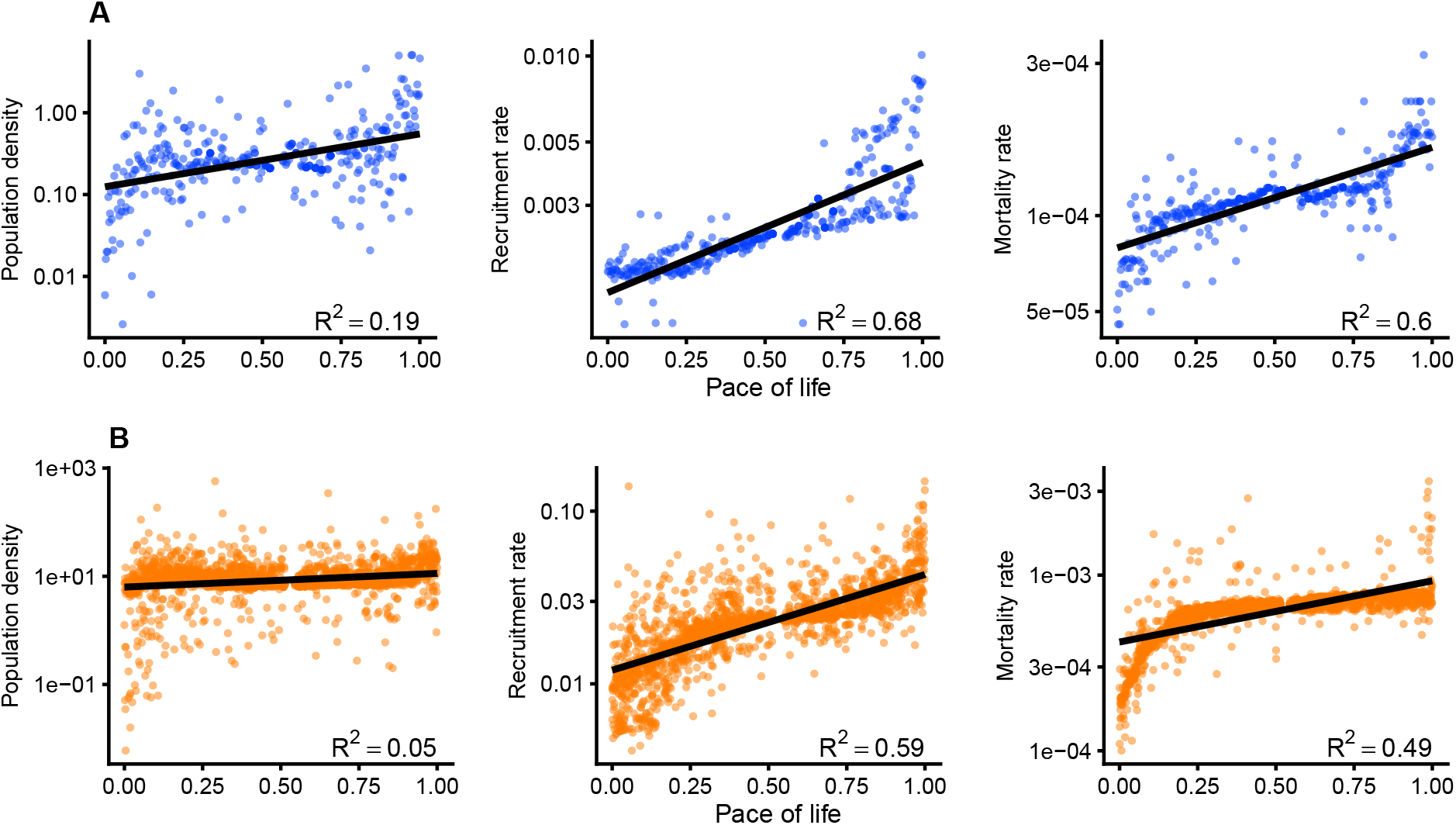
Fits of vertebrate host life history parameters to the pace of life measure (black lines) for (A) primates and (B) rodents with points indicating trait values from the underlying data set in blue and orange, respectively. Note that the y-axis is displayed on a log_10_-scale. Inset shows the *R*^2^ for the linear model log(*z*) ∼ Pace of life for each trait, *z*. The results shown here use generation time, the ratio of log(2) and the intrinsic population growth rate, to measure pace of life. The results for all other pace of life measures are given in the Supplementary Materials.

### Relating transmission potential to pace of life

The basic reproduction number, ℛ_0_, can be expressed as a function of the pace of life measure (*p*) to determine when ℛ_0_(*p*) is increasing. We evaluated a related quantity, the logarithmic derivative of ℛ_0_(*p*), 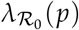, whose equation is given in (2). When 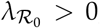, transmission potential increases along the life history continuum. 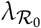 is itself a function of five quantities: the ratio of the blood meal demand and blood meal supply (*B*) as well as the logarithmic derivatives of the vertebrate host population density (*c*_*K*_), mortality rate (*c*_*µ*_), infection duration (*c*_*α*_), and susceptibility (*c*_*β*_), all of which may depend on *p*.

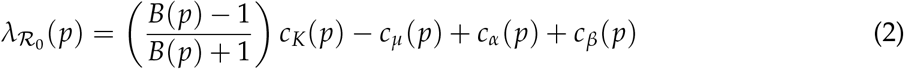

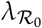 increases linearly with the magnitude of variation of immunological traits (*c*_*α*_ and *c*_*β*_). Therefore, greater variation in immunological traits can lead to 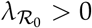 irrespective of any traits related to the mosquito or vertebrate host population density or mortality.

The dependence of 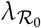 on contact dynamics between the vector and host is facilitated through the ratio of the total contact rate of the mosquito population if host availability was unlimited and the total contact rate with mosquitoes that the host population will tolerate (*B* = *σ*_*V*_*V*/*σ*_*H*_ *H*); *B* therefore quantifies the balance between the demand of the mosquito population for blood meals and the limited tolerance (or supply) of the host population to provide those blood meals (Thongsripong et al., 2021). If host tolerance is assumed to be unlimited, as is commonly assumed in modeling, then *B* = 0 and 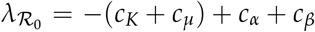. In this case, 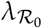 does not depend on any mosquito traits.

The nature of the relationship between 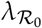 and *B* is determined by the sign of *c*_*K*_. If it is known whether population density increases (*c*_*K*_ > 0), decreases (*c*_*K*_ < 0), or is independent (*c*_*K*_ = 0) of pace of life, we can determine the intervals of *B* within which 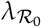 is positive. This, in turn, allows us to find bounds on the vector-host ratio, the ratio of vector abundance to host abundance, for which transmission potential increases along the life history continuum. One of the endpoints of these intervals is always either zero or infinity while the other is determined by the remaining parameters in equation (2).

### When does ℛ_0_ increase with pace of life?

After parameterizing the model, the values of 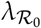 were calculated across all values of the pace of life measure *p*, parasite types, mosquito species, and temperatures. Figure 3 shows the range of values of 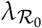 across all of these axes of variation. In almost every case, transmission potential increases along the life history continuum, i.e. 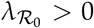. Only when the parasite causes chronic infections and all hosts are highly susceptible to infection do we see the possibility of 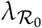 becoming negative. The estimates of 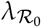 derived from the Ross-Macdonald model are represented by points which correspond with the minimum value of each of these ranges (Figure 3, black circles). While the range of 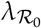 does vary across mosquito species, the minimum value is determined entirely by the parasite type. The ranges of 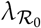 for lower level of immunological trait variation and for rodents are shown in Supplementary Figures S5 and S6.

**Figure 3:**
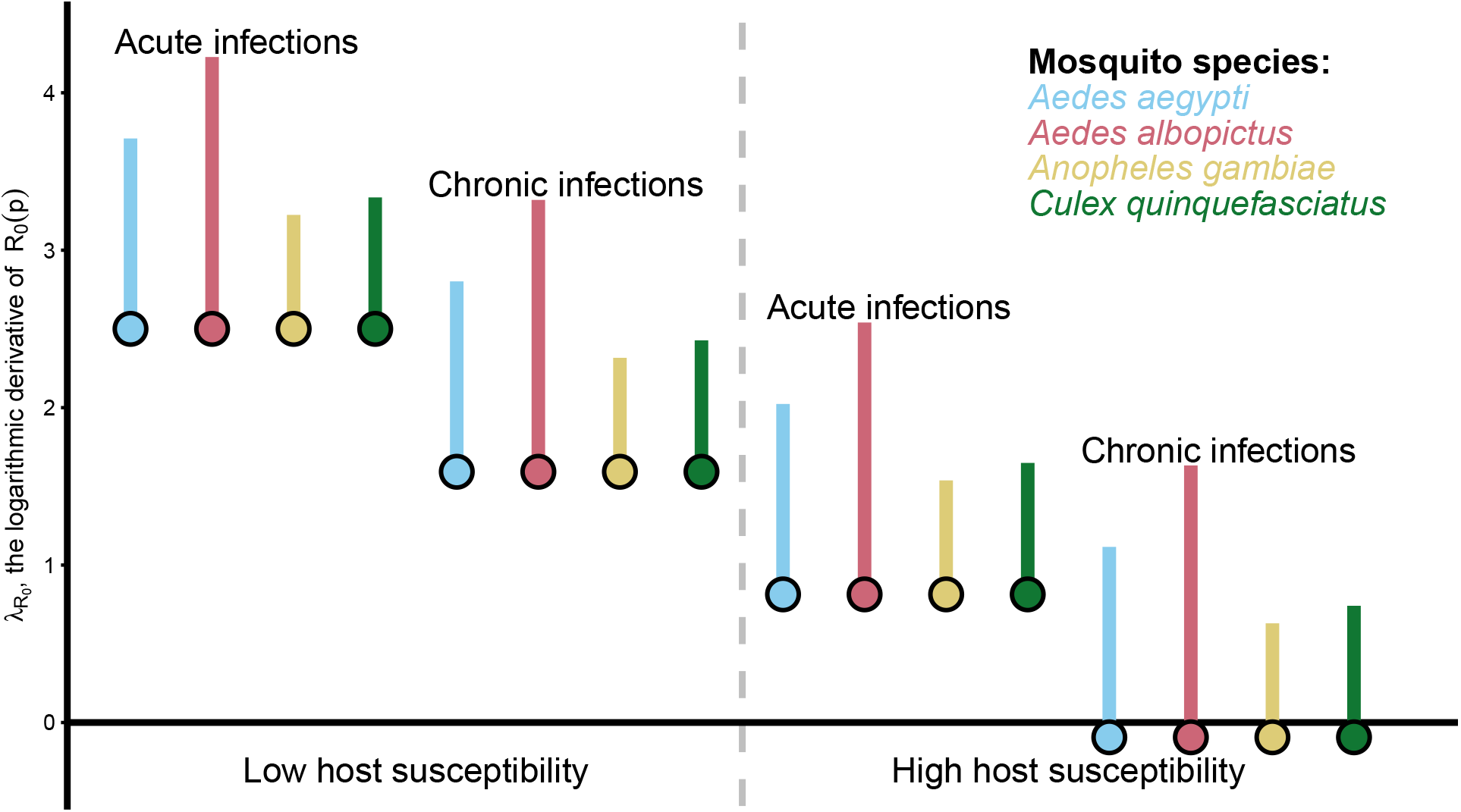
Ranges of 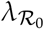 (see equation (2)) across the entire fast-slow life history continuum for each pathogen type and mosquito species as temperature is varied. Transmission potential increases with pace of life when 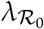 is positive. The horizontal axis distinguishes groupings of parasite type and mosquito species; it does not represent a variable.

### Critical vector-host ratio thresholds

In all cases, there is a minimum value of the vector-host ratio above which transmission potential always increases along the life history continuum. In fact, this minimum value was zero in all cases except when the parasite caused chronic infections and when hosts were highly susceptible to infection. In this case, we calculated the minimum vector-host abundance ratio at which 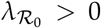 (Table 2) for two levels of immunological trait variation. When immunological trait variation was halved from wide to narrow, the lower bound for the vector-host ratio increased by an order of magnitude.

**Table 2:** Lower bounds for the vector-host ratio needed to guarantee that transmission potential increases along with pace of life. Values are shown for the case where vertebrate hosts are highly susceptible to parasites which induce chronic infections. All other parasite types had a lower bound of zero for the critical vector-host ratio. Higher levels of immunological trait variation lead to values for the critical vector-host ratio which are approximately one order of magnitude smaller.

| Mosquito species | Immunological trait variation | Vector-host ratio lower bound |
| --- | --- | --- |
| *Aedes aegypti* | Narrow | 100 |
|  | Wide | 8.49 |
| *Aedes albopictus* | Narrow | 99.6 |
|  | Wide | 8.41 |
| *Anopheles gambiae* | Narrow | 81.0 |
|  | Wide | 6.84 |
| *Culex quinquefasciatus* | Narrow | 109 |
|  | Wide | 9.21 |

## Discussion

To explore whether the transmission potential for mosquito-borne parasites increases with pace of life, we parameterized a mechanistic transmission model with trait data sets across several axes of variation including four mosquito vector species, two clades of vertebrate hosts, four immune response types, three levels of immunological trait variation, and a range of temperatures. Our results provide support for the pace of life hypothesis across all axes of variation except in the case of parasites causing chronic infections to which the host is highly susceptible. In this particular case, 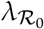 is still positive when the vector-host ratio is high (Table 2), which is often assumed to be the case in natural and human-dominated landscapes (although estimates of total adult mosquito population sizes are uncommon (Bowman et al., 2014; Silver, 2008)). These findings suggest that slower-lived hosts are more suitable reservoirs for mosquito-borne parasites in limited situations where i) the parasites cause long-lasting infections, ii) hosts are highly susceptible to infection, iii) mosquitoes are scarce or hosts are abundant or very tolerant of mosquito biting, and iv) the variation of immunological traits along the life history continuum is high. These restrictive requirements support the conjecture that we should expect fast-lived hosts to be more suitable reservoirs for mosquito-borne disease than slow-lived hosts.

These results were highly sensitive to the magnitude of variation in immunological traits because 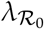 increases linearly with the amount of variation in infection duration and host susceptibility. When we explored the same analyses with narrower ranges of immune variation, we found that values of 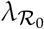 were lower than when immune traits varied more broadly. In fact, if immunological traits did not co-vary with pace of life, 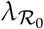 was always negative. Our findings further emphasize the need to quantify the relationship between immune function and life history variation to reliably determine the influence of pace of life on the parasite transmission potential of host populations, echoing the calls of other authors (Albery and Becker, 2021; Becker et al., 2019; Brace et al., 2017; Hawley and Altizer, 2011).

One strength of our approach is that we rely only on the amount of variation in immune traits and not their actual values. However, it is still not straightforward to determine to what extent immune traits vary across the pace of life spectrum of a given clade of host species. On its face, it might seem that immune function should always be strongly correlated with life history traits: longer-lived animals benefit significantly more from investment in immune function if they have more encounters with a wider variety of pathogens over their longer lifespans than short-lived animals (Valenzuela-Sánchez et al., 2021). At the same time, immune functions are costly, leading to trade-offs with growth and reproduction, particularly for short-lived animals (Brace et al., 2017). But these links between immune function and life history can be swamped by environmental effects, which can be a much stronger driver of immunological trait variation, particularly in birds and invertebrates (Sandland and Minchella, 2003; Tieleman, 2018).

The determinants of 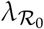 are the magnitudes of variation of the vertebrate host population density, mortality rate, infection duration, and susceptibility across trait space as well as the supply and demand of blood meals for the mosquito population. Population density, which determines the availability of hosts for blood-feeding by mosquitoes, provides the critical link between mosquito traits and transmission potential. When host availability did not limit mosquito contact rates – as is commonly assumed in modeling (Reiner et al., 2013) – the relationship between pace of life and transmission potential did not depend at all on the traits of the mosquito. Similarly, if population density of the vertebrate host was assumed to be independent of its pace of life (i.e. *c*_*K*_ = 0 in equation (2)), mosquito traits no longer affected 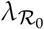. Whether population density is in fact a significant covariate of pace of life is a subject of debate (Manlove et al., 2022) and in our study population density exhibited the weakest correlation among all life history traits. While it seems reasonable to suggest that animals that successfully reproduce more rapidly will also form more dense populations, ecological trade-offs such as density-dependent mortality may act to restrict overall population sizes and decouple the relationship between reproduction and population density, particularly for contact-dependent transmission processes (Ostfeld et al., 2014; Valenzuela-Sánchez et al., 2021).

If population density and pace of life are indeed strongly linked, the direction of this relationship could determine whether ℛ_0_ increases with pace of life or if it decreases. For example, if population density does increase with pace of life, as was the case for the data used in our study (albeit weakly), then the vector-host ratio must be sufficiently large for ℛ_0_ to increase along with pace of life as shown in Table 2. If instead the population density of animals decreased with their pace of life, then these lower bounds transform into upper bounds, implying that transmission potential can only increase with pace of life when mosquito densities are much lower than that of vertebrate hosts.

Several factors excluded from this study might impact our findings. The movement patterns of hosts and vectors are important aspects of transmission and intrinsically linked to population densities across the landscape (Freed and Cann, 2013; Manlove et al., 2022). Previous modeling has shown that the dispersal patterns of hosts and vectors are key drivers of transmission, particularly when hosts disperse further than vectors or follow structured movement networks (Acevedo et al., 2015; Adams and Kapan, 2009; Prosper et al., 2012; Saucedo and Tien, 2022). A better understanding of how movement patterns of both vectors and vertebrates interact and influence the transmission dynamics of vector-borne diseases is needed before investigating the link between these patterns and life history and immunological traits.

In this study, we assumed that there was no intraspecific variation in life history or immune traits. However, the fitness effects of parasitic infection are key inter- and intra-specific drivers of immune trait variation (Becker et al., 2019). If parasite virulence affects host mortality or fecundity significantly, infection may itself confound the link between the pace of life and life history traits. Exposure to a novel parasite could lead to dramatic declines in host populations, further decoupling population density from life history (Daszak et al., 2003; Samuel et al., 2015). Improving our understanding of the links between immunological and life history traits would enable a more complete accounting of how variations in life history and immune traits affect the influence of the pace of life on transmission potential.

In light of the increased emergence rate of zoonoses and their incredible impact on human populations across the globe, identifying those species which pose the greatest risk of harboring and spreading zoonoses is a critical scientific task. Global surveillance of all potential reservoirs is not feasible, especially considering that not all existing possible zoonotic pathogens have been identified. Ecological theory can point to aspects of host populations which make them more likely to harbor and transmit zoonoses. The pace of life hypothesis of disease ecology, one among many theories suggesting which species should be targeted for surveillance, suggests that those species that “live fast and die young” are more likely be reservoirs for disease. Our findings provide evidence for this hypothesis for mosquito-borne parasites, as long as variations in immune function and pace of life are linked in hosts. Ultimately, comparative studies of host susceptibility and infection duration would provide the most valuable insights into what types of animals present the highest risk of harboring these pathogens.

## Supporting information

Supplement: Mathematical Details

## Acknowledgments

This work was supported by the NSF Ecology and Evolution of Infectious Diseases program (DEB 1717282 to BAH, SO, JPS, JD). The authors would also like to thank Éric Marty for valuable feedback and assistance with preparing the figures in the manuscript.

## Statement of Authorship

**Kyle Dahlin:** Conceptualization, Methodology, Software, Formal analysis, Data Curation, Writing - Original Draft, Writing - Review & Editing, Visualization, **Suzanne O’Regan:** Conceptualization, Methodology, Writing - Review & Editing, Supervision, Funding acquisition, **John Paul Schmidt:** Conceptualization, Methodology, Writing - Review & Editing, Funding acquisition, **Barbara Han:** Conceptualization, Writing - Review & Editing, Funding acquisition, **John Drake:** Conceptualization, Writing - Review & Editing, Supervision, Project administration, Funding acquisition

## Data and Code Availability

Data and code that support the ndings of this study are available in a GitHub repository (https://github.com/DrakeLab/dahlin-paceoflife-mbps).

