## Supplementary Figures for "Fast-lived Vertebrate Hosts Exhibit Higher Potential for Mosquito-borne Parasite Transmission"

Kyle J.-M. Dahlin[1,2*]

Suzanne M. O'Regan[1,2]

John Paul Schmidt[1,2]

Barbara A. Han[3]

John M. Drake[1,2]

1. Odum School of Ecology, University of Georgia, Athens, Georgia 30602, USA;

2. Center for the Ecology of Infectious Diseases, University of Georgia, Athens, Georgia 30602, USA;

3. Cary Institute of Ecosystem Studies, Millbrook, New York 12545, USA.

### Supplementary Figures and Tables

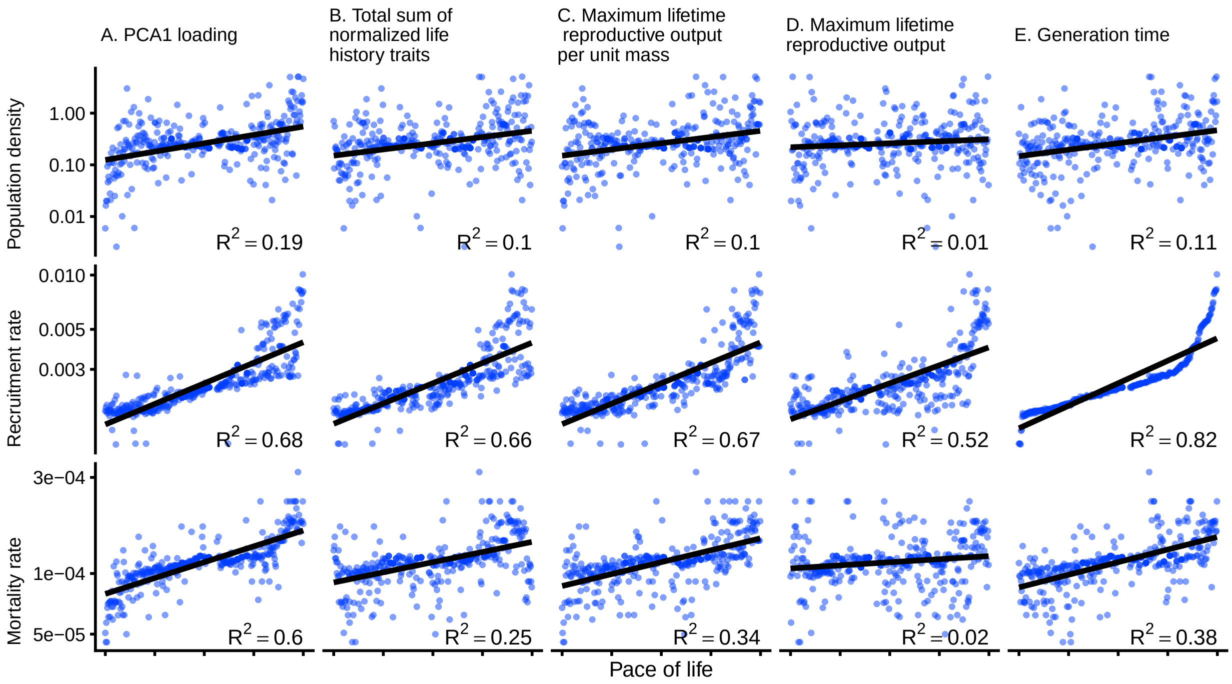


Figure S1: Fits of vertebrate host life history parameters to each candidate pace of life measure for primates. Blue points indicate values taken from the trait data set. Note that the y-axis is displayed on a $\log_{10}$-scale. Inset shows the $R^2$ for the linear model $\log(z) \sim$ Pace of life for each trait, $z$. Generation time (E.) was chosen as it best approximated the covariation among the life history traits. The full definition of each candidate is given in the mathematical supplement.

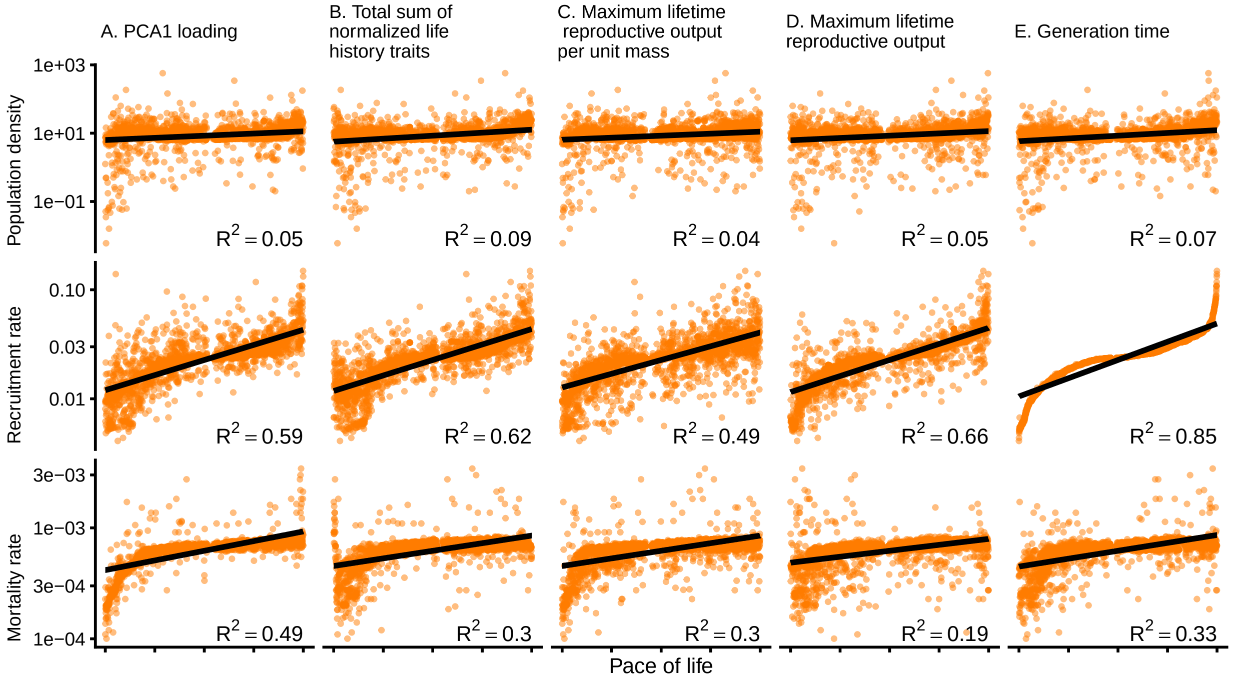


Figure S2: Fits of vertebrate host life history parameters to each candidate pace of life measure for rodents. Orange points indicate values taken from the trait data set. Note that the y-axis is displayed on a $\log_{10}$-scale. Inset shows the $R^2$ for the linear model $\log(z) \sim$ Pace of life for each trait, $z$. Generation time (E.) was chosen as it best approximated the covariation among the life history traits. The full definition of each candidate is given in the mathematical supplement and results for the other candidate pace of life metrics are shown in Supplementary Figures S1 and S2.

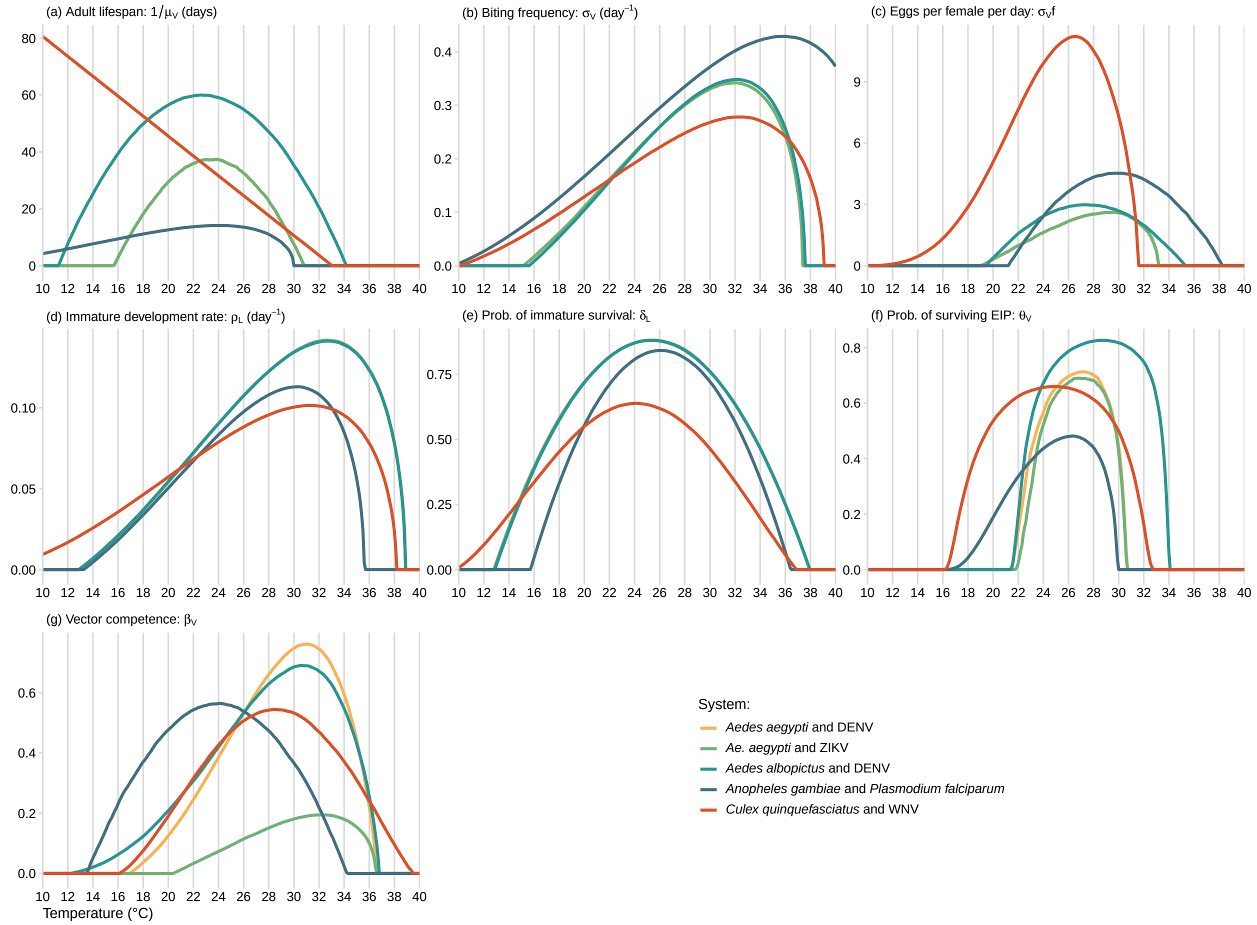
(a) Adult lifespan: $1/\mu_V$ (days)
(b) Biting frequency: $\sigma_V$ (day$^{-1}$)
(c) Eggs per female per day: $\sigma_V f$
(d) Immature development rate: $\rho_L$ (day$^{-1}$)
(e) Prob. of immature survival: $\delta_L$
(f) Prob. of surviving EIP: $\theta_V$
(g) Vector competence: $\beta_V$
Temperature (°C)
System:
Aedes aegypti and DENV
Ae. aegypti and ZIKV
Aedes albopictus and DENV
Anopheles gambiae and Plasmodium falciparum
Culex quinquefasciatus and WNV

Figure S3:

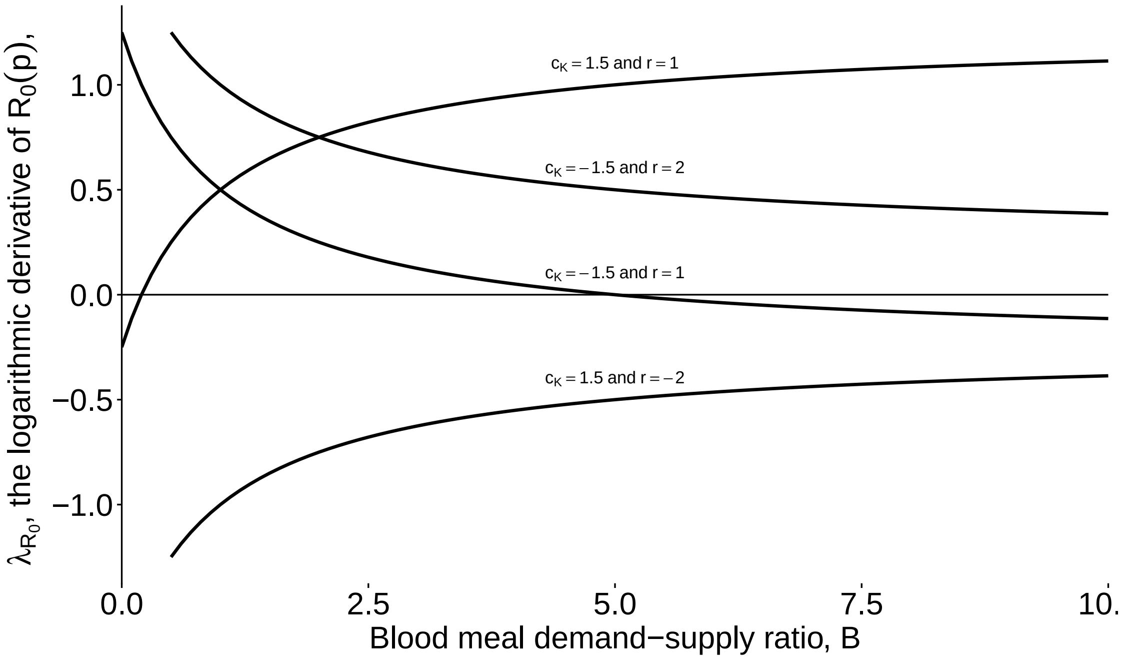


Figure S4: Representative curves showing the relationship between $\lambda_{\mathcal{R}_0}$ (equation (1) in the main text) and the blood meal demand-supply ratio, $B$. The pace of life hypothesis suggests that the curve should always lie above zero (implying that transmission potential always increases with the pace of life history of the host). $c_K$ is the logarithmic derivative of vertebrate host population density with respect to pace of life history while $r$ is a relative measure of the variation with respect to pace of life history of immune traits (infection duration and susceptibility) compared to mortality rate. When $c_K$ is negative, the curve slopes downwards and, if $r$ is also sufficiently large, $\lambda_{\mathcal{R}_0}$ is always positive (otherwise it undergoes a sign change). On the other hand, when $c_K$ is positive, the curve slopes upwards and a sufficiently large r value is necessary for $\lambda_{\mathcal{R}_0}$ to be positive.

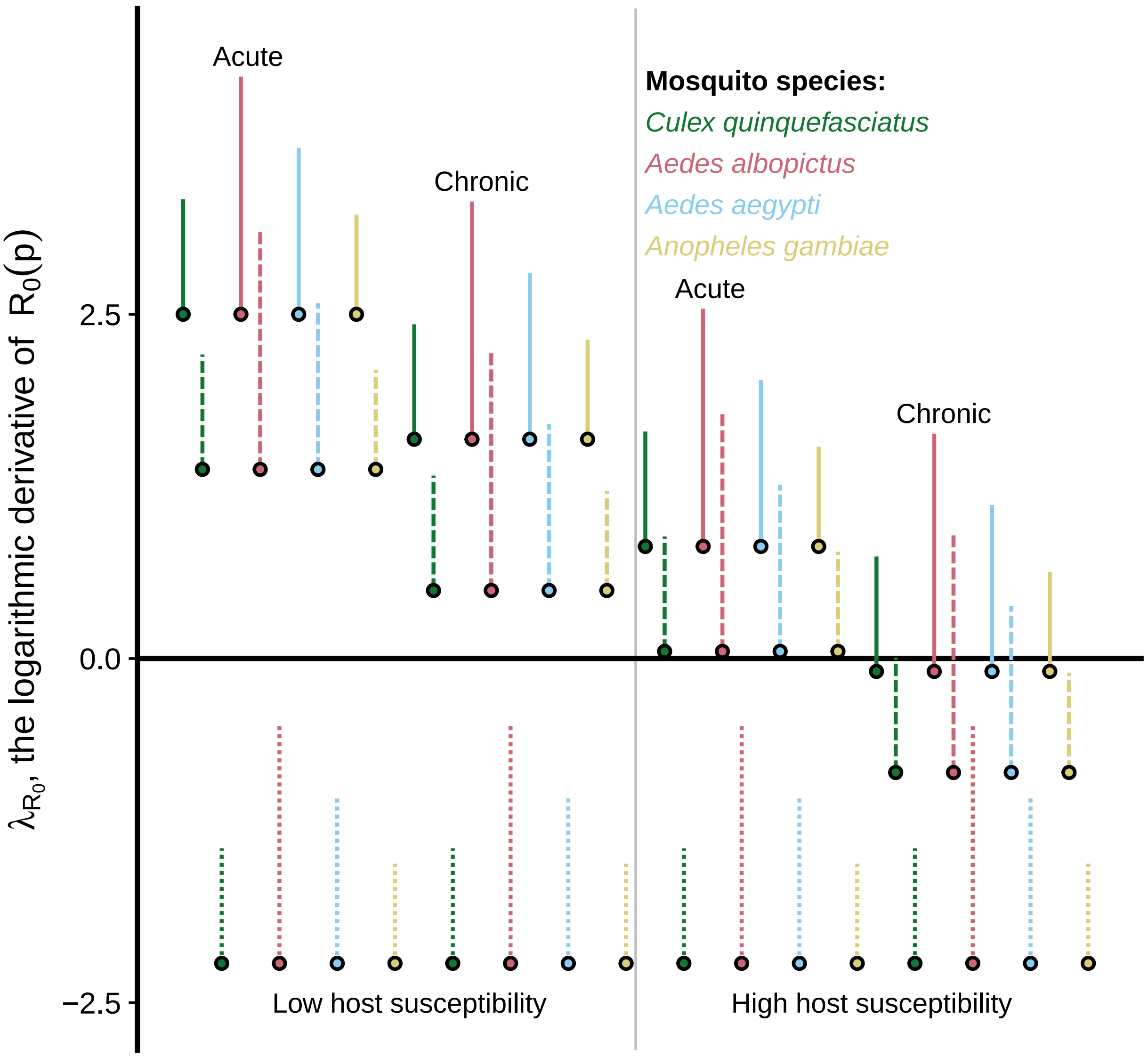


Figure S5: Ranges of values of $\lambda_{\mathcal{R}_0}$ for primate traits. The level of opacity indicates the strength of the relationship between immunological traits (susceptibility and infection duration) and pace of life history. The most transparent bars indicate no relationship between immunological traits and pace of life history. The other levels of variation are described in Table 1 of the main text.

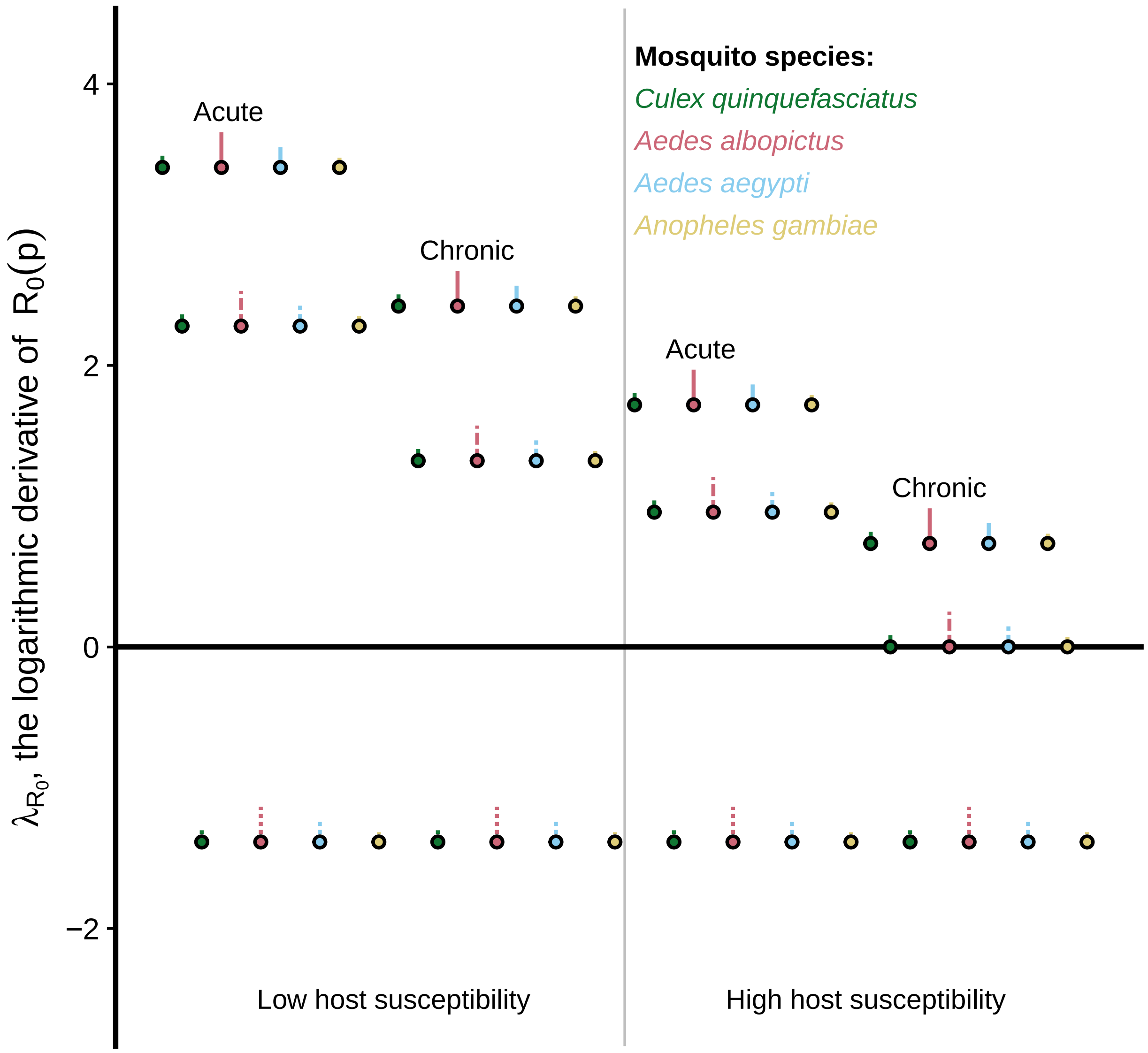


Figure S6: Ranges of values of $\lambda_{\mathcal{R}_0}$ for rodent traits. The level of opacity indicates the strength of the relationship between immunological traits (susceptibility and infection duration) and pace of life history. The most transparent bars indicate no relationship between immunological traits and pace of life history. The other levels of variation are described in Table 1 of the main text.
