## Supplement: Mathematical Details for "Fast-lived Vertebrate Hosts Exhibit Higher Potential for Mosquito-borne Parasite Transmission"

### S1. Parameterizing host parameters

We aim to represent the three life history parameters: recruitment rate ($\lambda_H$), mortality rate ($\mu_H$), and carrying capacity ($K_H$) as continuous functions of a pace of life parameter ($p$). The life history parameters (simulated values) are related to life history traits (real or imputed data derived from PANTHERIA) as follows: $\lambda_H$ as the product of litter size ("15-1_LitterSize") and litters per day ("16-1_LittersPerYear"), $\mu$ as the inverse of maximum longevity ("17-1_MaxLongevity_m"), and $K_H$ as population density ("21-1_PopulationDensity_n/km2"). All naming conventions in quotes are derived from the PANTHERIA data set. Unit conversions were made to ensure that rates are in units of per day and density in per hectare.

The life history parameters are fit to pace of life in three steps. First several candidate metrics for pace of life are calculated from the trait data set. Then each life history parameter is fit as a function of each candidate metric. Finally, the metric which best represents the covariation of life history traits with pace of life, as measured by the sum of squares of $R^2$ values, is chosen.

S1.1. **Quantifying pace of life.** There is no single commonly-accepted method of quantifying pace of life commonly accepted [4]. We therefore considered several candidate metrics for pace of life and evaluated which best described the observed trait data. We require only one condition on these metrics: that recruitment and mortality rates should both increase with pace of life.

The first metric we considered was the **first principal component** derived from the full data set, excluding non-numeric traits and age at eye opening ("2-1_AgeatEyeOpening_d") and body length of weanlings ("13-3_WeaningHeadBodyLen_mm") as there were not enough data to impute these traits reliably. The loading of the first principal component was most strongly positively associated with population density (0.10), litter size (0.12), and litters per year (0.20), and strongly negatively associated with longevity (-0.20).

The remaining metrics are defined and calculated as described below.

**Sum of all life history traits considered in Van de Walle et al. [13]:**
1/(Age at first birth)+Longevity+Sexual maturity period/(Interbirth interval×Longevity)+Litters per day
- Each summand was normalized before calculation

**Maximum lifetime reproductive output:**
MLRO = LitterSize × SexualMaturityPeriod/InterbirthInterval

**Maximum lifetime reproductive output per unit mass:**
MLROperMass = MLRO/AdultMass

**Duration of lifespan spent in the sexually mature stage:**
SexualMaturityPeriod = Longevity − AgeOfSexualMaturity

**Generation time (or population doubling time):**
log(2)/(LittersPerDay × LitterSize − 1/Longevity)

[1]Odum School of Ecology, University of Georgia 140 E. Green Street, Athens, GA 30602, USA
[2]Center for the Ecology of Infectious Diseases, University of Georgia 203 D.W. Brooks Drive, Athens, GA 30602, USA
[3]Cary Institute of Ecosystem Studies Box AB, Millbrook, NY 12545, USA

Each metric was calculated for each species in the data set, then these values were normalized to the interval between zero and one using the function `percent_rank` from the R package dplyr. In cases where the directionality of the relationship between a metric and traits needed to be specified (i.e. in the principal component analysis), the direction was set so that recruitment and mortality rates increased along with the pace of life metric. Supplementary Figures S1 and S2 exhibit the relationship between each pace of life metric and life history traits from the data set (colored points).

S1.2. **Life history parameters.** The life history parameters were modeled as exponential functions of pace of life: $z(p) = z_0 e^{cp}$. We also explored modeling the parameters as linear functions of $p$ ($z(p) = z_0 + cp$) but this resulted in models which more poorly explained the covariation between $p$ and the parameters than the exponential model while having the same number of parameters ($z_0$ and $p$). Supplementary Figures S1 and S2 show the fitted curves of each life history parameter as a function of pace of life (black lines) as well as the coefficient of determination. Generation time (A.) was chosen for $p$ because it had the highest sum of squares $R^2$ value across the three life history traits.

### S2. Description of the Compartmental Model

We use a mathematical model of the transmission of a mosquito-borne pathogen between populations of mosquitoes and sylvatic hosts as a system of ordinary differential equations originally presented in [3]. The state variables represent abundances of vertebrate hosts (denoted with a subscript $H$) and mosquitoes (subscript $V$). The vertebrate host and mosquito populations are each divided into susceptible ($S_x$) and infectious ($I_x$) compartments. The full system of equations is given by (S2.1):

$$
\begin{aligned}
\frac{dS_H}{dt} &= \lambda_H H - \left(\frac{\lambda_H - \mu_H}{K_H}\right) H^2 - b_H(H,V)\,\beta_H \frac{I_V}{V} S_H - \mu_H S_H,\\
\frac{dI_H}{dt} &= b_H(H,V)\,\beta_H \frac{I_V}{V} S_H - (\gamma_H + \mu_H) I_H,\\
\frac{dR_H}{dt} &= \gamma_H I_H - \mu_H R_H,\\
\frac{dS_V}{dt} &= \lambda_V - b_V(H,V)\,\beta_V \frac{I_H}{H} S_V - \mu_V S_V,\\
\frac{dI_V}{dt} &= b_V(H,V)\,\beta_V \theta_V \frac{I_H}{H} S_V - \mu_V I_V.. 
\end{aligned}
\tag{S2.1}
$$

$H = S_H + I_H + R_H$ represents the total vertebrate host abundance and $V = S_V + I_V$ the total mosquito abundance. The functions $b_H$ and $b_V$ represent the density-dependent contact rates between hosts and vectors, described in S2.1. Model parameters and their definitions are given in Table 1 of the main text and in Table S1.

Vertebrate host population dynamics follow a logistic model. The mosquito population dynamics are described by a first-order linear approximation of a logistic model (constant recruitment and constant per-capita mortality rates). A sub-model of the population dynamics of the aquatic stage for mosquitoes (described fully in Section S2.2) is used to determine the recruitment rate for adult mosquitoes, $\lambda_V$. We assume that mosquito reproduction is not dependent on the presence of the vertebrate host.

The pathogen is assumed to be transmitted solely through the mosquito vector. Hosts recover from infection at the rate $\gamma_H$ and attain lifelong immunity. After exposure to the pathogen, mosquitoes must survive the extrinsic incubation period of the pathogen in order to become infectious (represented by the term $\theta_V$). Mosquitoes remain infectious for the remainder of their lifespan.

S2.1. **Contact rates.** The host biting rate, $b_H$, is the number of mosquito bites received per unit time per vertebrate host. The mosquito biting rate, $b_V$, is the number of blood meals taken from vertebrate hosts per unit time per mosquito. We consider two formulations for these contact rates: the commonly-used Ross-MacDonald contact rate model (abbreviated RM) [9] and the Chitnis dynamic contact rate model [2]. Both contact rate models are parametrized by the parameters $\sigma_H$ and $\sigma_V$, which are the maximum per-host and per-mosquito interspecific contact rates, respectively. The parameter $\sigma_V$ can be measured empirically in a laboratory setting (for example, by allowing colonized female mosquitoes to feed as frequently as they like) or estimated through measurement of the length of the mosquito gonotrophic cycle. On the other hand, $\sigma_H$, the maximum number of mosquito bites an individual host is willing to tolerate per unit time, is much more difficult to measure. For brevity, we will refer to $\sigma_H$ as the biting tolerance of the vertebrate host.

The Chitnis dynamic contact rate (originally presented in [2]) models host behavioral defenses and intolerance to mosquito biting by incorporating density dependence in both contact rates. In the Chitnis model, the total number of mosquito-vertebrate host contacts per unit time, $b$, is estimated as half the harmonic mean of the supply of blood meals ($\sigma_H H$) and the demand for blood meals ($\sigma_V V$).

$$b\left(H,V\right)=\frac{\sigma_H H\sigma_V V}{\sigma_H H+\sigma_V V} \tag{S2.2}$$

Then $b_H$ ($b_V$) is obtained by dividing $b$ by the total host population $H$ (the total mosquito population $V$):

$$b_H\left(H,V\right)=\sigma_H\left(\frac{\sigma_V V}{\sigma_H H+\sigma_V V}\right), \tag{S2.3}$$

$$b_V\left(H,V\right)=\sigma_V\left(\frac{\sigma_H H}{\sigma_H H+\sigma_V V}\right). \tag{S2.4}$$

The standard Ross-Macdonald contact rate formulation can then be obtained from the Chitnis formulation by taking the limit as biting tolerance, $\sigma_H$, approaches infinity:

$$b_H^{\mathrm{RM}}\left(H,V\right)=\lim_{\sigma_H\to\infty}b_H\left(H,V\right)=\sigma_V\left(\frac{V}{H}\right), \tag{S2.5}$$

$$b_V^{\mathrm{RM}}\left(H,V\right)=\lim_{\sigma_H\to\infty}b_V\left(H,V\right)=\sigma_V. \tag{S2.6}$$

In the Ross-MacDonald model, the host biting rate, $b_H$, is a product of the maximum mosquito biting rate and the vector-host ratio. The mosquito biting rate is simply the maximum mosquito biting rate, $\sigma_V$. Thus, the primary difference between the Chitnis dynamic contact rate model and the Ross-MacDonald contact rate model is the existence of biting tolerance thresholds ($\sigma_H$) in the Chitnis model.

We assume that mosquito reproduction is not limited by the abundance of the focal host. Typically, mosquitoes which pose a zoonotic risk are somewhat generalist in their feeding: although they may specialize on feeding on mammals or birds, for example, they often feed on multiple species within these classes [8, 11]. The feeding choice of these mosquitoes is often driven more so by host abundance than by the particular preference of the mosquito species [1]. These mosquitoes are thus capable of opportunistically utilizing alternate blood meal sources when one source is rare.

S2.2. **Immature (aquatic-stage) mosquito population dynamics.** The recruitment rate of immature mosquitoes is assumed to be proportional to both the biting frequency of adult mosquitoes and the total abundance of adult mosquitoes. Immature mosquitoes are also assumed to have a density-dependent mortality rate, inducing a limited carrying capacity of larval mosquitoes (as is observed across many species).

Let $L$ represent the population of immature mosquitoes, $K_L$ the carrying capacity of immature mosquitoes, and $\mu_L$ the mortality rate of immature mosquitoes. Then the dynamics of the entire mosquito population, in the absence of pathogen transmission, can be represented by the system:

$$\frac{dL}{dt}=\sigma_V f\left(1-\frac{L}{K_L}\right)V-\left(\rho_L+\mu_L\right)L, \tag{S2.7}$$

$$\frac{dV}{dt}=\rho_L L-\mu_V V. \tag{S2.8}$$

This system has two equilibria: $(L_0,V_0)=(0,0)$ and $(L^*,V^*)$ where $L^*=K_L\left(1-\frac{\mu_V}{\sigma_V f\delta_L}\right)$ and $V^*=\frac{\rho_L L_0}{\mu_V}$. Here, $\delta_L=\rho_L/\left(\rho_L+\mu_L\right)$ represents the probability that an immature mosquito survives to emergence as an adult.

| Parameter | Our symbol | *Aedes aegypti* | *Aedes albopictus* | *Anopheles* spp. | *Culex quinquefasciatus* |
|---|---|---|---|---|---|
| Biting rate | $\sigma_V$ | $a$ | $a$ | $a$ | $a$ |
| Fecundity | $f$ | $0.5 \times \frac{\text{EFD}}{a}$ | $0.5 \times \frac{\text{EFD}}{a}$ | $0.5 \times \frac{\text{EFD}}{a}$ | $0.5 \times \frac{\text{ER}\times p_O}{a}$ |
| Pr(egg to adult survival) | $\delta_L$ | $p_{EA}$ | $p_{EA}$ | $p_{EA}$ | $\text{EV} \times p_{LA}$ |
| Development rate | $\rho_L$ | MDR | MDR | MDR | MDR |
| Adult mortality rate | $\mu_V$ | $\frac{1}{\text{lf}}$ | $\frac{1}{\text{lf}}$ | $-\frac{1}{\log(p)}$ | $\frac{1}{\text{lf}}$ |
| Vector competence | $\beta_V$ | $bc$ | $bc$ | $bc$ | $bc$ |
| Original source | | [5, 6, 12] | [5, 6] | [5, 7] | [5, 10] |

Table S1. The thermal performance curves for the parameters used in (S2.1) are derived from trait thermal performance curves presented in past works, listed under original source. In the original sources, $a$ is the (constant) mosquito biting rate, EFD is eggs per female per day, $p_O$ is the probability of oviposition, $p_{\text{EA}}$ is the probability of an egg surviving to adulthood, $EV$ is egg viability, $p_{\text{LA}}$ is the probability of a larval mosquito surviving to adulthood, MDR is the mosquito development rate, lf is adult mosquito lifespan, and $p$ is the daily probability of survival for an adult mosquito.

Henceforth, we assume that the immature mosquito population is at equilibrium. Then the autonomous (but temperature-dependent) adult recruitment rate is given by

$$\lambda_V = \rho_L L^* = \rho_L K_L \left(1 - \frac{\mu_V}{\sigma_V f \delta_L}\right) \tag{S2.9}$$

S2.3. **Temperature-dependent mosquito traits.** In our model, mosquito parameters are determined from thermal performance curves (TPCs) originally presented in [5, 7, 10]. Each trait thermal performance curve is either a linear, quadratic, or Briere function of temperature. To translate the notation used in previous works to match with the parameters of our model, several substitutions were necessary. In [7], quadratic functions of temperature are written as $Q_1(T) = q_1 T^2 + rT + s$ while in [5] and [10], the function is written as $Q_2(T) = -q_2 (T - T_{\min})(T - T_{\max})$. To translate the parameters of [7] into the thermal performance curves parameters used in [5, 10], we used the following transformations:

$$q_2 = -q_1, \tag{S2.10}$$

$$T_{\min} = \frac{-r + \sqrt{r^2 - 4q_1 s}}{2q_1}, \tag{S2.11}$$

$$T_{\max} = \frac{-r - \sqrt{r^2 - 4q_1 s}}{2q_1}. \tag{S2.12}$$

Mosquito parameters in (S2.1) and (S2.7) were determined from values previously reported in other sources. Table S1 show how our model parameters are derived from previously reported thermal performance curves and the source from which they were obtained.

### S3. Equilibrium Analysis of the Model

In this Section, we determine the equilibria of (S2.1)and calculate the basic and type reproduction numbers of (S2.1).

S3.1. **Infection-free equilibria.** The model admits equilibria in the absence of active parasite transmission both with and without hosts or mosquitoes. Assuming no parasites in the system, the system (S2.1) reduces to (S3.2)

$$\frac{dH}{dt} = rH\left(1 - \frac{H}{K_H}\right), \tag{S3.1}$$

$$\frac{dV}{dt} = \lambda_V - \mu_V V, \tag{S3.2}$$

where $r = \lambda_H - \mu_H$. Henceforth, we define $H^* = K_H$ and $V^* = \frac{\lambda_V}{\mu_V}$. We assume that $r$ is positive, meaning that the host population size increases when it is low. Then, in the absence of pathogens, there are four equilibria: $E^* = (K_H, 0, 0, V^*, 0)$, $E_H = (K_H, 0, 0, 0, 0)$, $E_V = (0, 0, 0, V^*, 0)$, and $E_0 = (0, 0, 0, 0, 0)$. We are interested in the case where both species populations are persistent ($E^*$). Hence, we refer to $E^*$ as the disease-free equilibrium or DFE. In the absence of the pathogen, $E^*$ is locally asymptotically stable if $r > 0$ and $\lambda_V > 0$. Note that the disease-free equilibrium mosquito population size is a function of temperature through the two parameters $\lambda_V$ and $\mu_V$.

Henceforth, we assume that the DFE is stable in the absence of pathogens ($r > 0$ and $\lambda_V > 0$) and that total host and mosquito abundances are at their DFE values ($H = K_H$ and $V = V^*$). In this case, we can make the substitutions: $S_H = K_H - I_H - R_H$ and $S_V = V^* - I_V$ to arrive at a reduced version of (S2.1):

$$\begin{aligned}
\frac{dI_H}{dt} &= b_H^*\left(\beta_H \frac{I_V}{V^*}\right)(K_H - I_H - R_H) - (\gamma_H + \mu_H) I_H, \\
\frac{dR_H}{dt} &= \gamma_H I_H - \mu_H R_H, \\
\frac{dI_V}{dt} &= b_V^*\left(\beta_V \frac{I_H}{K_H}\right)\theta_V (V^* - I_V) - \mu_V I_V.
\end{aligned} \tag{S3.3}$$

Here we have used the shorthand $b_H^* = b_H(K_H, V^*)$ and $b_V^* = b_V(K_H, V^*)$ to more simply refer to the contact rates at the DFE.

S3.2. $\mathcal{R}_0$ **and the type reproduction numbers.** We calculate the basic reproduction number using the method of van den Driessche and Watmough [14] to obtain the next generation matrix

$$\boldsymbol{K} = \begin{bmatrix} 0 & b_H^*\left(\beta_H \frac{K_H}{V^*}\right)/\mu_V \\ b_V^*\left(\beta_V \frac{V^*}{K_H}\right)\theta_V/(\gamma_H + \mu_H) & 0 \end{bmatrix} \tag{S3.4}$$

and the basic reproduction number

$$\mathcal{R}_0 = \sqrt{\left(b_V^* \beta_V \theta_V \frac{1}{\gamma_H + \mu_H}\right)\left(b_H^* \beta_H \frac{1}{\mu_V}\right)}. \tag{S3.5}$$

$\mathcal{R}_0$ can be decomposed into host-to-vector ($\mathcal{R}_H$) and vector-to-host ($\mathcal{R}_V$) the type reproduction numbers

$$\mathcal{R}_H = b_V^*\left(\beta_V \frac{V^*}{K_H}\right)\theta_V\left(\frac{1}{\gamma_H + \mu_H}\right), \tag{S3.6}$$

$$\mathcal{R}_V = b_H^*\left(\beta_H \frac{K_H}{V^*}\right)\left(\frac{1}{\mu_V}\right). \tag{S3.7}$$
